# Local differentiation in the defensive morphology of an invasive zooplankton species is not genetically based

**DOI:** 10.1101/098707

**Authors:** Giuseppe E. Fiorino, Andrew G. McAdam

**Affiliations:** Department of Integrative Biology, University of Guelph, 50 Stone Road East, Guelph, ON, Canada, N1G 2W1

**Keywords:** *Bythotrephes longimanus*, cladoceran, common garden experiment, invasive species, local adaptation, phenotypic plasticity

## Abstract

Evolutionary changes in functional traits represent one possible reason why exotic species spread to become invasive, but empirical studies of the mechanisms driving phenotypic differentiation between populations of invasive species are rare. This study tested whether differences in distal spine length among populations of the invasive cladoceran, *Bythotrephes longimanus*, could be explained by local adaptation or phenotypic plasticity. We collected *Bythotrephes* from six lakes and found that distal spine lengths and natural selection on distal spine length differed among populations, but were unrelated to the gape-limitation of the dominant fish predator in the lake from which they were collected. A common garden experiment revealed significant genetic and maternal variation for distal spine length, but phenotypic differences among populations were not genetically based. Phenotypic differences among lakes in this ecologically important trait are, therefore, the result of plasticity and not local adaptation, despite spatially variable selection on this heritable trait. The ability of *Bythotrephes* to plastically adjust distal spine length may explain the success of this species at invading lake ecosystems with diverse biotic environments.

## Introduction

Invasive species have substantial adverse impacts on global biodiversity, community structure, and ecosystem function (Vitousek et al. 1996; Mack et al. 2000), but few exotic species spread, causing the wide-scale ecological and economic damage that we associate with “invasiveness” (Mooney and Cleland 2001). Understanding why some exotic species become invasive while others do not has been the focus of decades of ecological and ecosystem-level research (Drake et al. 1989; Novak 2007). More recently, there has been increasing interest in the role of evolutionary changes in biological invasions (Mooney and Cleland 2001; Lee 2002; Parker et al. 2003; Lambrinos 2004; Facon et al. 2006), but tests for adaptive evolution remain rare.

The spread of an exotic species depends on its ability to perform well in new biotic and abiotic conditions (Shea and Chesson 2002; Facon et al. 2006). One way in which this could be achieved is through local adaptation (Lee 2002; Parker et al. 2003; Lambrinos 2004; Facon et al. 2006), which is the process whereby divergent natural selection (i.e. selection that differs among habitats) causes populations to become genetically differentiated (Kawecki and Ebert 2004). Natural selection might be particularly strong in exotic species because species introduced into a foreign environment often encounter new resources, competitors, or predators (Mooney and Cleland 2001; Lambrinos 2004). Additionally, for populations to locally adapt, there must be sufficient genetic variation underlying the traits experiencing divergent selection (Lynch and Walsh 1998). For exotic species, invasions characterized by large founder populations or a large number of founder events are expected to have high genetic variance, whereas invasions characterized by small founder populations or a small number of founder events are expected to have reduced genetic variance, which may constrain local adaptation (Allendorf and Lundquist 2003; Lockwood et al. 2005).

Local adaptation results in phenotypic differentiation between populations, but the presence of such differentiation is not sufficient to demonstrate that populations are locally adapted. Phenotypic plasticity is the ability of a genotype to produce alternate phenotypes based on environmental conditions (Pigliucci 2005), and represents an alternative mechanism by which exotic species can adaptively respond to heterogeneity in their environment. Plasticity can, therefore, produce a pattern of phenotypic differentiation among populations that is consistent with that of local adaptation, but without the underlying genetic differences (Kawecki and Ebert 2004). Additionally, because plasticity allows different genotypes to produce the same phenotype, it can reduce the strength of selection, which might constrain local adaptation (Pfennig et al. 2010). Alternatively, plasticity can allow organisms to cope with new environments where they might not otherwise persist, which can result in novel selection pressures that subsequently cause local adaptation (West-Eberhard 2003).

Despite the potential importance of local adaptation and plasticity to invasiveness (Parker et al. 2003), little is known about their relative importance with respect to the spread of exotic species. Most previous work has focused on invasive plants, where limited studies suggest that local adaptation and phenotypic plasticity are not mutually exclusive. For example, Si et al. (2014) found evidence for local adaptation and phenotypic plasticity of several growth characteristics that contributed to the successful invasion of *Wedelia trilobata* across a tropical island. Similarly, Godoy et al. (2011) found that local adaptation and phenotypic plasticity were both involved in the successful invasion of the heavily shaded understory of South American evergreen temperate rainforest by *Prunella vulgaris* (also see: Parker et al. 2003). Dybdahl and Kane (2005) provided a rare example outside of plants, in which North American populations of invasive freshwater snails (*Potamopyrgus antipodarum*) were found to show phenotypic plasticity (but not local adaptation) for life history and growth traits that facilitated their spread. In order to better understand the general mechanisms by which exotic species spread (and hence, become invasive), further empirical studies of the mechanisms driving phenotypic differentiation between populations of invasive species for ecologically important traits are needed.

The spiny water flea, *Bythotrephes longimanus* (hereafter, *Bythotrephes*), is an invasive species in the Laurentian Great Lakes and many surrounding inland lakes where it negatively impacts lake ecosystems due to its central position in the food web as a predator of zooplankton (Bunnell et al. 2011) and prey for fish (Pothoven et al. 2007). The tail spine of *Bythotrephes* is used as a morphological defense against fish predation (Barnhisel 1991a, b), and previous work on five Canadian Shield lakes identified that *Bythotrephes* in lakes dominated by gape-limited fish predators experience natural selection for longer distal spines (i.e. the posterior-most segment of the tail spine), whereas those in lakes dominated by non-gape-limited predators experience no selection on distal spine length (Miehls et al. 2014). Gape-limited predation (GLP) occurs when predators cannot consume individuals of a focal prey species above a certain size determined by the gape-size of the predator, and is generally expected to cause natural selection for increased size in prey (Day et al. 2002; Urban 2008). In contrast, non-gape-limited predation (NGLP), in which predators are not constrained by mouth size, is expected to impose no selection on the size of prey (Urban 2007, 2008). Additionally, distal spine length of *Bythotrephes* from Lake Michigan is heritable (*H^2^* = 0.27-0.76; Miehls et al. 2012), and differences in mean distal spine length among Canadian Shield lakes were consistent with differences in natural selection among lakes: *Bythotrephes* from lakes dominated by GLP were found to have 17% longer distal spines compared to lakes dominated by NGLP (Miehls et al. 2014). These findings suggest that local adaptation might explain the observed phenotypic differentiation among populations of *Bythotrephes*. However, cladocerans are commonly phenotypically plastic, especially for traits involved in predator defense. For example, the cladoceran *Daphnia lumholtzi* produces neonates with longer head spines when exposed to fish predator kairomones (Dzialowski et al. 2003) (also see: Lüning 1992). This maternal induction of offspring phenotypes in response to the maternal environment is called a maternal effect (Mousseau and Fox 1998). It has also been shown that *Bythotrephes* from Lake Michigan induce longer distal spines in offspring in response to warmer water temperature (but not fish kairomones), which may act as an indirect cue for natural selection associated with GLP (Miehls et al. 2013). It is thus plausible that local differences in distal spine length among populations of *Bythotrephes* could be explained by either local adaptation in response to GLP or maternal induction of longer distal spines in offspring in response to an environmental cue associated with GLP.

In this study, we first measured *Bythotrephes* distal spine lengths and the strength of natural selection on distal spine length in six Canadian Shield lakes that differed in the presence or absence of GLP. We then conducted a common garden experiment (Kawecki and Ebert 2004) to evaluate the hypotheses of local adaptation and phenotypic plasticity as causes of phenotypic differences among these populations. Using clonal lines (Lynch and Walsh 1998), we reared individuals from the six study lakes in identical conditions for two generations. We measured genetic and maternal variation for distal spine length to determine broad-sense heritability for the trait and maternal effects, and determined whether phenotypic differences in distal spine length among populations were genetically based. If phenotypic differences among populations were due to local adaptation (i.e. genetically based differentiation), we predicted that phenotypic differences in distal spine length among populations would be maintained in second-generation *Bythotrephes* reared in a common garden environment. Alternatively, if phenotypic differences among populations were due to plasticity, we predicted that phenotypic differences among populations would no longer be present in second generation *Bythotrephes*.

## Material and Methods

### Study Species

*Bythotrephes* is a predatory cladoceran zooplankter with a widespread native distribution throughout the Palearctic region (Therriault et al. 2002; Colautti et al. 2005; Kim and Yan 2013), which can tolerate a wide range of pH, salinity, temperature, and conductivity (Grigorovich et al. 1998). *Bythotrephes* was first identified in the Laurentian Great Lakes in the early 1980s (Johannsson et al. 1991), and has since spread to more than 160 inland Ontario lakes, and lakes in the mid-western USA (Kelly et al. 2013). In the Canadian Shield, *Bythotrephes* can be found in lakes dominated by gape-limited predators, such as rainbow smelt (*Osmerus mordax*), or non-gape limited predators, such as cisco (*Coregonus artedi*) (Strecker et al. 2006; Young and Yan 2008). Rainbow smelt are considered gape-limited predators of *Bythotrephes* because they start consuming these prey at ~10 cm in length (Barnhisel and Harvey 1995), but are not significant predators of *Bythotrephes* as adults (Young and Yan 2008). Alternatively, cisco are considered non-gape-limited predators because they are significant predators of *Bythotrephes* as adult fish (Young and Yan 2008).

The *Bythotrephes* caudal process (i.e. tail spine) consists of segments, the longest of which is the distal spine (i.e. the section from the posterior tip of the spine to the first paired articular spines; Fig. 1), which is present at birth and does not change in length with development (Burkhardt 1994). Thus, the length of the distal spine cannot respond plastically to the environment directly experienced by offspring, but it can be maternally induced (Miehls et al. 2013). Only the distal spine is present in neonates (i.e. the first instar stage), but total spine length increases through development when an additional spine segment is added to the base of the spine at each instar molt (Branstrator 2005). These segments are each separated by paired articular spines (Fig. 1), which allow the instar stage to be identified and the length of each segment to be measured separately (Yurista 1992). Like most cladocerans, *Bythotrephes* have a cyclically parthenogenetic life cycle, reproducing apomictically (i.e. clonally) multiple times before reproducing sexually at the end of the growing season (Yurista 1992; Branstrator 2005). Apomictic reproduction produces eggs that immediately develop into young in the brood pouch, whereas sexual reproduction results in resting eggs that overwinter on the lake bottom before hatching the following spring (Yurista 1992; Branstrator 2005).

**Fig. 1.**
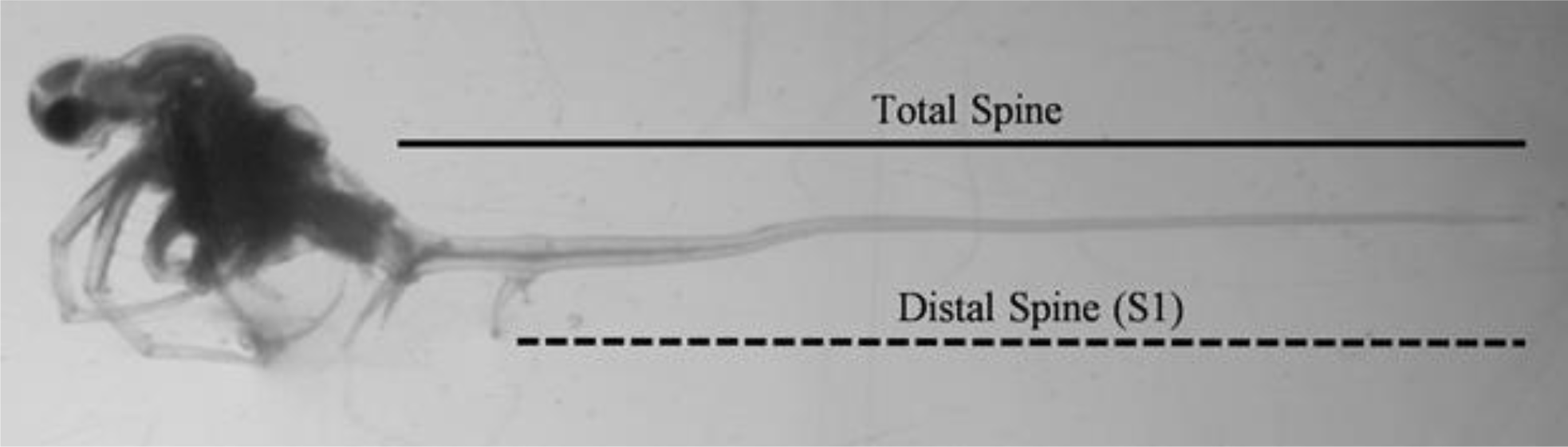
Photograph of *Bythotrephes longimanus*. The total tail spine (solid line) is composed of several segments. The distal spine segment (i.e. the section from the posterior tip of the spine to the first paired articular spines) is present at birth and does not grow. Total spine length increases during development through the production of additional spine segments at each instar molt. The photographed individual can be identified as a second instar animal because it has two spine segments separated by two pairs of articular spines.

(Graphic created using Adobe Photoshop CC 2015.1)

### Study Lakes

*Bythotrephes* were collected from six lakes in the Muskoka district and County of Haliburton in south-central Ontario (Online Resource, Fig. S1). Predation on *Bythotrephes* in three of the lakes (Peninsula, Mary, and Fairy; hereafter, GLP lakes) is thought to be dominated by the gape-limited fish predator, rainbow smelt, while in the three other lakes (Boshkung, Harp, and Drag; hereafter, NGLP lakes) predation is thought to be dominated by the non-gape-limited fish predator, cisco (Strecker et al. 2006; Young and Yan 2008; S. J. Sandstrom and N. Lester, unpublished data). Although rainbow smelt are present in Boshkung Lake (Young and Yan 2008), cisco have been reported to be more abundant and were considered to be the dominant *Bythotrephes* predator (Strecker et al. 2006; Miehls et al. 2014), so Boshkung was classified as a NGLP lake (Table 1).

**Table 1.** Field information, dominant predation regime, categorical predator abundance, and water temperature at the time of sampling for each lake from which *Bythotrephes longimanus* were sampled.

| Lake | Sample Date | GPS Coordinates | Dom. Pred. | Smelt Abund. | Cisco Abund. | Temp. (°C) |
| --- | --- | --- | --- | --- | --- | --- |
| Peninsula | July 28, 2014 | 45°21.1'N, 79°06.1'W | GLP | M | A | 21.5 |
| Mary | July 29, 2014 | 45°13.5'N, 79°16.2'W | GLP | M | A | 21.2 |
| Fairy | July 29, 2014 | 45°19.4'N, 79°11.3'W | GLP | M | A | 21.7 |
| Boshkung | July 30, 2014 | 45°03.1'N, 78°43.3'W | NGLP | L | M | 20.5 |
| Harp | July 28, 2014 | 45°22.5'N, 79°08.1'W | NGLP | A | H | 21.0 |
| Drag | July 30, 2014 | 45°05.2'N, 78°24.2'W | NGLP | N/A | N/A | 20.5 |

GLP = lake dominated by gape-limited predation (i.e. rainbow smelt); NGLP = lake dominated by non-gape-limited predation (i.e. cisco). Categorical fish abundances were not available for Drag Lake. Water temperature was taken 1 m below the surface immediately prior to sampling.
Sources: S. J. Sandstrom and N. Lester, Ontario Ministry of Natural Resources (unpublished data), Strecker et al. (2006), Young and Yan (2008).

These lakes were chosen because they were invaded by *Bythotrephes* over the last 30 years and are characteristic of many lakes in the Canadian Shield that are dominated by rainbow smelt or cisco (Strecker et al. 2006; Young and Yan 2008). They are similar to one another physically (in terms of depth and water temperature) and biologically (in terms of invertebrate predators, and other fish predators of *Bythotrephes*) (Hovius et al. 2006; Strecker et al. 2006; Young and Yan 2008). Although Harp Lake is much smaller than the other lakes, several studies have found that it is ecologically similar (Hovius et al. 2006; Strecker et al. 2006; Young and Yan 2008). Five of these six lakes (excluding Drag) were previously used by Miehls et al. (2014) to test for the effects of GLP on natural selection and local differences in distal spine length. Miehls et al. (2014) sampled an additional lake (Kashagawigamog, NGLP) but sample sizes were too low to measure natural selection. We sampled Drag Lake (NGLP) instead of Kashagawigamog to balance the experimental design.

### Sample Collection

*Bythotrephes* were collected between 10 am and 2 pm over three days during the middle of the growing season (July 29-31, 2014) using a conical zooplankton net with a 0.5 m diameter opening and 363 μm mesh size. To measure distal spine lengths and natural selection for each lake, *Bythotrephes* were collected with the net horizontally towed at a depth of 10-15 m, and approximately 100 individuals were haphazardly chosen and immediately preserved in 95% ethanol. For the common garden experiment, live *Bythotrephes* were collected using a vertical net tow (instead of a horizontal tow) through the top 15 m of the water column, and 30-40 actively swimming individuals without pigmented brood pouches were individually isolated in 60 mL jars containing 50 mL of lake water filtered through a 63 μm sieve. A vertical net tow was used to collect individuals for the common garden experiment because it minimizes damage to the animals associated with turbulence, whereas a horizontal tow was used to collect individuals to measure phenotypic differences and selection because animals could be collected in greater quantity. Additionally, a vertical tow was used to account for potential differences in *Bythotrephes* diel vertical migration among lakes. Diel vertical migration by *Bythotrephes* is greater in lakes dominated by rainbow smelt (GLP lakes) compared to those dominated by cisco (NGLP lakes) (Young and Yan 2008). However, in all our study lakes, individuals are not commonly found below 15 m (Young and Yan 2008), meaning a vertical net tow of 15 m provided a representative sample of all *Bythotrephes* in the water column, regardless of which lake was sampled. Furthermore, we ensured that there was no difference in *Bythotrephes* distal spine length between the two different towing methods (Online Resource, Appendix B). All collection methods were based on those reported by Kim and Yan (2010) and Miehls et al. (2014).

### Common Garden Experiment

We used a common garden experiment and clonal breeding design (Fig. 2) to measure genetic and maternal variation for distal spine length, and to determine whether phenotypic differences among populations were genetically based. During the three-day collection period, all live *Bythotrephes* were maintained at the Dorset Environment Science Center (DESC, Dorset, Ontario) in a climate controlled facility (20°C, 14L:10D photoperiod) in lake water filtered through a 63 μm sieve from their “home” lake. Afterwards, the cultures were moved to an environmental chamber at the Hagen Aqualab (University of Guelph, Guelph, Ontario) under the aforementioned temperature and photoperiod and introduced to the common garden medium. The medium was an autoclaved mixture of lake water filtered through a 63 μm sieve from the six sampled lakes (i.e. each *Bythotrephes* was reared in water that was 1/6^th^ of their local environment). *Bythotrephes* received daily water changes and were fed *ad lib* with approximately 150 *Artemia* sp. nauplii that were less than 30 h old (Miehls et al. 2012).

**Fig. 2.**
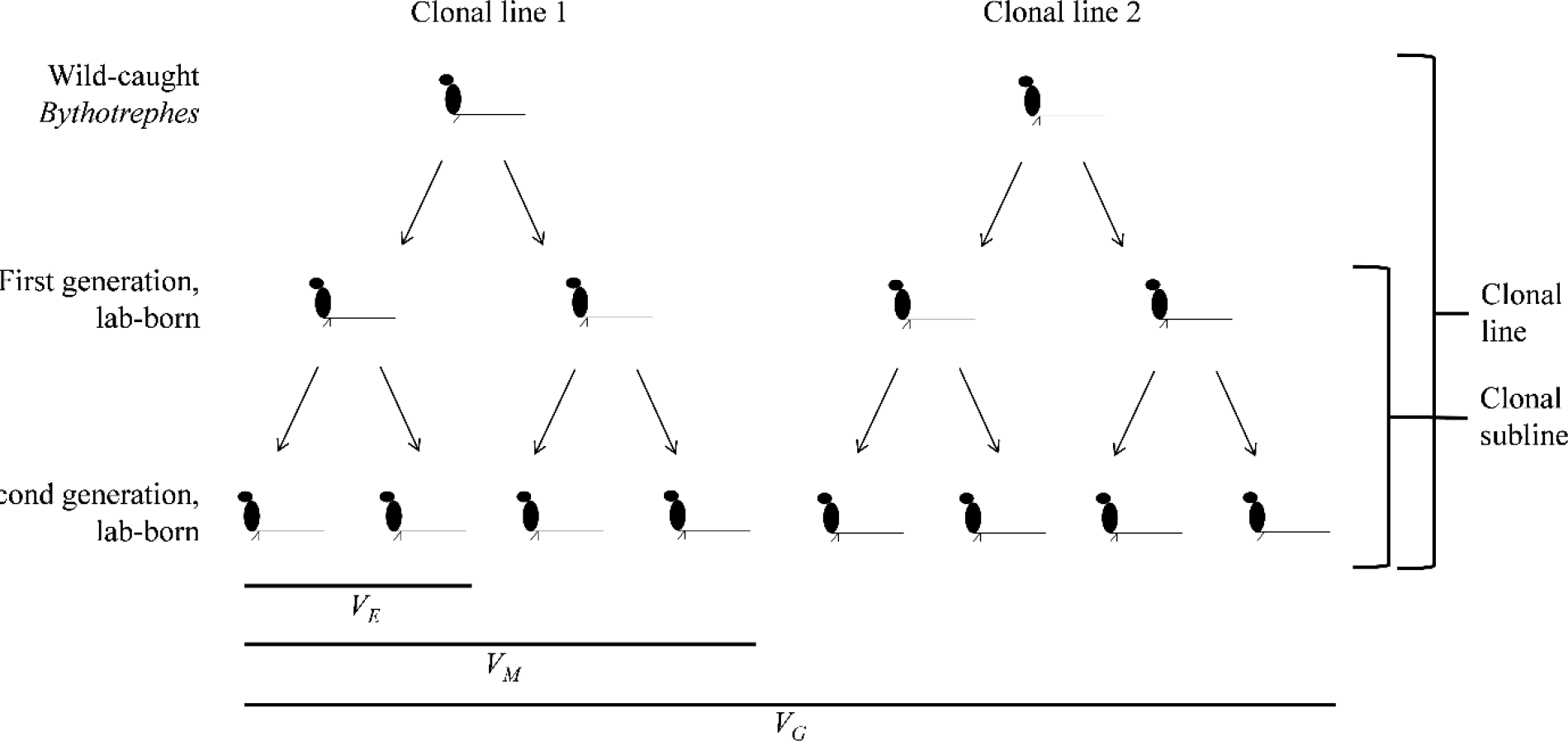
Schematic diagram of *Bythotrephes longimanus* clonal analysis design. Wild-caught individuals were used to initiate clonal lines. All offspring from wild-caught individuals (first generation lab-born) were used to initiate clonal sublines. Distal spine length measurements of second-generation lab-born animals were analyzed to estimate variance components and to determine if populations were genetically differentiated. The variance in distal spine length among clonal lines represents the genetic variance (*V_G_*), the variance among sublines within clonal lines represents maternal variance (*V_M_*), and the variance among individuals within clonal sublines is the environmental variance (*V_E_*). This figure is modified from Lynch and Walsh (1998) and Miehls et al. (2012).

(Graphic created using Microsoft PowerPoint 2013)

Clonal lines were initiated using 188 wild-caught individuals (28-37 per lake), and were reared in the common garden through two apomictic generations (Fig. 2; Miehls et al. 2012, 2013). Once a female produced offspring she was preserved in 95% ethanol within 24 h. All offspring were individually transferred to 60 mL jars containing 50 mL of common garden medium, also within 24 h (Miehls et al. 2012). Of the 188 clonal lines, 12.2% produced second-generation lab-born offspring (7-37 individuals per lake; Online Resource, Table S1). We conducted a supplementary analysis to ensure that lab mortality did not bias our results (Online Resource, Appendix C).

### Measurement

All *Bythotrephes* were photographed using a digital camera mounted to a dissecting microscope (Leica MZ8, Leica Microsystems). IMAGEJ software (Abramoff et al. 2004) was used to measure the length of the distal spine segment from the tip of the tail spine to the first paired articular spines (Fig. 1) to the nearest 0.001 mm. Instar was assessed by counting the number of paired articular spines on the total tail spine (Fig. 1).

### Comparing Distal Spine Length among Natural Populations

We tested for phenotypic differences in mean distal spine length among lake populations using one-way ANOVA with distal spine length of wild-caught, first instar individuals as the response variable and lake as the predictor, where a significant effect of lake would indicate that mean distal spine length differed among lakes. For this model, Tukey’s multiple comparison test (Abdi and Williams 2010) was used to assess the significance of differences among pairs of lakes. To determine whether there was phenotypic differentiation between predation regimes (i.e. GLP vs. NGLP), we fitted a linear mixed-effects (LME) model using the *nlme* package in R (Pinheiro et al. 2015) with the distal spine length of wild-caught, first instar individuals as the response variable, predation regime as a fixed effect, and lake as a random effect (to account for variation among lakes unrelated to predation type). In this model, a significant effect of predation regime would indicate that mean distal spine length was associated with the gape-limitation of the dominant fish predator. For both models, only first instar individuals were considered because distal spine length at this stage represents the pre-selection phenotype; therefore, using only first instar *Bythotrephes* ensured that differences in distal spine length among lakes were not confounded by selection.

### Measuring Natural Selection

Natural selection was quantified by comparing distal spine lengths between first and second instar *Bythotrephes* collected on the same day (i.e. from different cohorts). Because *Bythotrephes* distal spine length does not change with development (Burkhardt 1994), a difference in mean distal spine length between first and second instar individuals from different cohorts represents the relationship between distal spine length and survival between first and second instar stages, and not developmentally based differences. For example, an environment with strong gape-limited predation should favour first instar individuals with longer distal spines over those with shorter distal spines. As a result, we would expect second instar individuals from that cohort (i.e. post-selection individuals) to have longer distal spines, on average, than first instar individuals from a different cohort (i.e. pre-selection individuals). The magnitude and direction of the difference represents the strength and direction of natural selection (Miehls et al. 2014). Although *Bythotrephes* develops into a third or fourth instar stage, the comparison between the first two stages was assessed because the distal spine represents the entire length of the spine in first instar individuals, which was the expected target of selection (Miehls et al. 2014).

We calculated selection differentials (Falconer and Mackay 1996) for each population as the difference between the mean distal spine lengths for the first two instar stages (Miehls et al. 2014). The statistical significance of these selection differentials for each population was assessed using Welch *t*-tests (two-tailed). In these analyses, a statistically longer mean distal spine for second instar individuals compared to first instar individuals would indicate significant selection for longer distal spines in that lake. Additionally, to compare to other published estimates of selection, standardized selection differentials (i.e. selection intensities) were calculated by dividing the selection differential for a lake by the standard deviation of distal spine length for first and second instar animals from that lake (Miehls et al. 2014).

To statistically test whether selection on distal spine length differed among lakes, we used two-way ANOVA with distal spine length of wild-caught first and second instar animals as the response variable, lake and instar as predictors, and a lake-by-instar interaction. In this model, a significant effect of instar would indicate that there was selection on distal spine length irrespective of lake; a significant effect of lake would indicate that distal length differs by lake irrespective of selection; and a significant lake-by-instar interaction would indicate that selection differs among lakes. To determine whether selection varied consistently with predation regime, we fitted a LME model with distal spine length of wild-caught, first and second instar animals as the response variable, predation regime, instar, and a predation regime-by-instar interaction as fixed effects, and lake as a random effect. In this model, a significant effect of instar would indicate that there was selection on distal spine length irrespective of predation regime; a significant effect of predation regime would indicate that distal length differs by predation regime irrespective of selection; and a significant predation regime-by-instar interaction would indicate that selection depended on the gape limitation of the dominant fish predator.

### Determining Genetically Based Differences among Populations

We reared *Bythotrephes* from all study lakes in a laboratory setting under identical conditions for two generations to eliminate phenotypic differences among populations that may be expressed as a result of environmental heterogeneity among lake populations, including maternal effects (Mousseau and Fox 1998). As aforementioned, the *Bythotrephes* distal spine is present at birth and its length does not change with development (Burkhardt 1994). Therefore, the distal spine lengths of second-generation lab-born individuals are expressed in response to the common lab environment experienced by their mothers, and any remaining differences among populations should be genetically based (assuming negligible grand-maternal effects).

To statistically determine whether phenotypic differences in distal spine length among lakes were genetically based, we fitted a LME model with distal spine length of second-generation individuals as the response variable, lake as a fixed effect, and clonal subline nested within clonal line as random effects. In this model, a nonsignificant effect of lake would indicate that distal spine length was not genetically differentiated among lakes, which would be consistent with a phenotypic plasticity hypothesis. Alternatively, a significant effect of lake would indicate that local differences in distal spine length were genetically based, and could thus reflect local adaptation in response to GLP.

### Estimating Broad-sense Heritability and Maternal Effects

The clonal breeding design that we used (Fig. 2) allowed for the quantification of genetic (*V_G_*), maternal (*V_M_*), and environmental (*V_E_*) variance components for distal spine length (Lynch and Walsh 1998; Miehls et al. 2012). We fitted a LME model with distal spine length of second-generation lab-born individuals as the response variable, the intercept as the only fixed effect, and clonal subline nested within clonal line as random effects. In an additional model, lake was included as a fixed effect but this did not alter our conclusions (Online Resource, Table S2). In this breeding design, the variance in distal spine length among clonal lines estimates genetic variance, the variance among sublines within clonal lines estimates maternal variance, and the variance within sublines estimates environmental variance. Note, because *Bythotrephes* distal spine length is fixed from birth, the variance within sublines (i.e. environmental variance) must be due to small scale environmental differences within the brood pouch of the mother during development. Similarly, variance in distal spine length among sublines (i.e. maternal variance) could be confounded by environmental differences within the brood pouch of the grandmother during development of the mothers, or subtle differences experienced by the mothers in the lab. To assess the significance of the variance components, we obtained 95% confidence intervals around the random effects (Pinheiro and Bates 2000) and conducted model comparisons using likelihood ratio tests (see *Assessing the Significance of Random Effects*) (Miehls et al. 2014). We calculated broad-sense heritability (*H^2^*) as the ratio of among-line variance to the total phenotypic variance (i.e. the sum of the among-line, among-subline, and within-subline variances), and calculated maternal effects (*m^2^*) as the ratio of among-subline variance to the total phenotypic variance.

### Assessing the Significance of Random Effects

For all LME models, the statistical significance of the random effects was assessed through model comparisons using likelihood ratio tests in which the change in the log-likelihood between the more complex model and the simpler model was assumed to follow a chi-squared distribution where the degrees of freedom were equal to the difference in the number of parameters between the more complex and simpler models (df = 1 in all cases here). For models with one random effect, we fitted one additional model with the same fixed effects but without the random effect. The significance of the random effect was assessed by comparing these two models. For nested models (i.e. models with multiple nested random effects), we fitted additional models with the same fixed effects but with successively fewer random effects, starting with the removal of the most nested random effect. The significance of a random effect was assessed by comparing the model that included the random effect of interest to the simpler model without that random effect. All statistical analyses were conducted using R version 3.2.2 (R Core Team 2015). The statistical assumptions of homoscedasticity and normality were met for all models.

## Results

### Mean Distal Spine Lengths of Natural Populations

The mean distal spine length of all wild-caught, first instar individuals was 5.80 ± 0.02 mm (mean ± SE, SD = 0.33 mm). Mean distal spine length differed among lakes (ANOVA: *F_4,225_* = 17.1, *P* < 0.001), but did not differ by predation regime (Table 2; Fig. 3). Specifically, populations in Peninsula Lake (GLP) and Boshkung Lake (NGLP) had significantly longer distal spines than populations in Mary Lake (GLP), Fairy Lake (GLP) and Harp Lake (NGLP) (Tukey HSD: *P* < 0.003). There was no difference in mean distal spine length between the Peninsula and Boshkung populations (Tukey HSD: *P* = 0.913) and no differences among the Mary, Fairy, and Harp populations (Tukey HSD: *P* > 0.244). No first instar animals were collected from Drag Lake.

**Table 2.** The mean distal spine length of wild-caught *Bythotrephes longimanus* did not differ between predation regimes (i.e. GLP vs. NGLP).

| Fixed | *F* | df | *P* |
| --- | --- | --- | --- |
| Predation Regime | 0.001 | 1,3 | 0.979 |
| Random | $\sigma^2$ | $\chi^2_1$ | *P* |
| Lake | 0.06 | 37.3 | < **0.001** |

Results are reported for a linear mixed-effects model in which only the distal spine lengths of first instar individuals were included as the response variable. The significance of the lake random effect was assessed using a likelihood ratio test that compared the model to a model with the same fixed effect but with no random lake effect. *F* = *F*-statistic; df = degrees of freedom; *P* = *P*-value; $\sigma^2$ = among-lake variance; $\chi^2_1$ = Chi-square value with 1 degree of freedom.

**Fig. 3.**
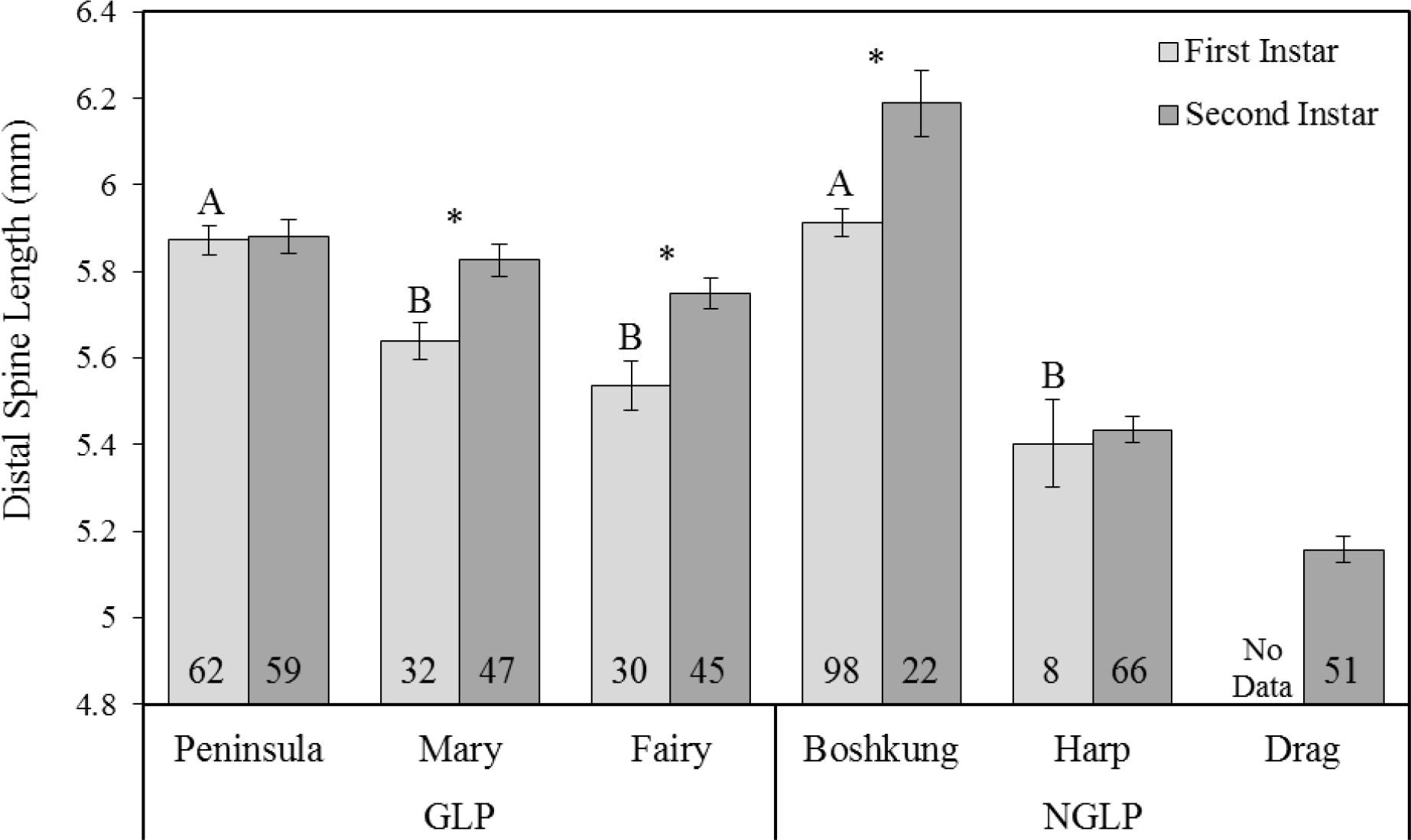
Mean distal spine length of first and second instar wild-caught *Bythotrephes longimanus* for all study lakes. Mean distal spine lengths for first instar animals labelled with different letters were significantly different from one another (Tukey HSD: *P* < 0.003). The difference in mean distal spine length between first and second instars represents the selection differential for that lake. Asterisks represent significant directional selection for increased distal spine length for that lake (Welch *t*-tests: *P* < 0.003; Table 3). No first instar animals were collected from Drag Lake. The number in each bar represents the sample size for that lake. Error bars represent ± 1 standard error.

(Graphic created using Microsoft Excel 2013)

### Natural Selection

Differences in distal spine length between first and second instar *Bythotrephes* differed by lake (i.e. a significant lake-by-instar interaction; ANOVA: *F_4,459_* = 3.5, *P* = 0.008; Fig. 3), indicating that strength of natural selection differed among populations in the study lakes. However, differences in selection among lakes were not consistently related to predation regime (i.e. a nonsignificant predation regime-by-instar interaction; Table 4). Of the GLP lakes, there was significant directional selection for increased distal spine length in Mary and Fairy (i.e. the mean distal spine length in second instar individuals was larger than that of first instar individuals), but selection on distal spine length in Peninsula was not significant. Of the NGLP lakes, selection was not significant in Harp, but there was significant directional selection for increased distal spine length in Boshkung (Table 3). Natural selection could not be assessed for Drag Lake because no first instar animals were collected from this lake.

**Table 3.** Natural selection on *Bythotrephes longimanus* distal spine length in lakes where fish predation was dominated by gape-limited predation (GLP) or non-gape-limited predators (NGLP).

| Predation | Lake | *t* | df | *P* | *S* | 95% CI of *S* |  | SD | *i* |
| --- | --- | --- | --- | --- | --- | --- | --- | --- | --- |
|  |  |  |  |  |  | Lower | Upper |  |  |
| GLP | Peninsula | 0.17 | 116 | 0.868 | 0.01 | -0.09 | 0.11 | 0.28 | 0.03 |
|  | Mary | 3.31 | 68 | **0.002** | 0.19 | 0.07 | 0.30 | 0.26 | 0.71 |
|  | Fairy | 3.15 | 51 | **0.003** | 0.21 | 0.08 | 0.35 | 0.29 | 0.74 |
| NGLP | Boshkung | 3.33 | 28 | **0.002** | 0.28 | 0.11 | 0.45 | 0.34 | 0.82 |
|  | Harp | 0.30 | 8 | 0.769 | 0.03 | -0.21 | 0.27 | 0.24 | 0.13 |

Selection was measured as the difference in mean distal spine length between second instar animals and first instar animals for each population. Selection for Drag Lake could not be calculated because no first instar animals were collected. *t* and *P* are derived from t-tests comparing distal spine length of first and second instar animals. *t* = *t*-statistic; df = degrees of freedom; *P* = *P*-value; *S* = selection differential; CI = confidence interval; SD = pooled standard deviation for first and second instar individuals; *i* = standardized selection differential (i.e. selection intensity).

**Table 4.**
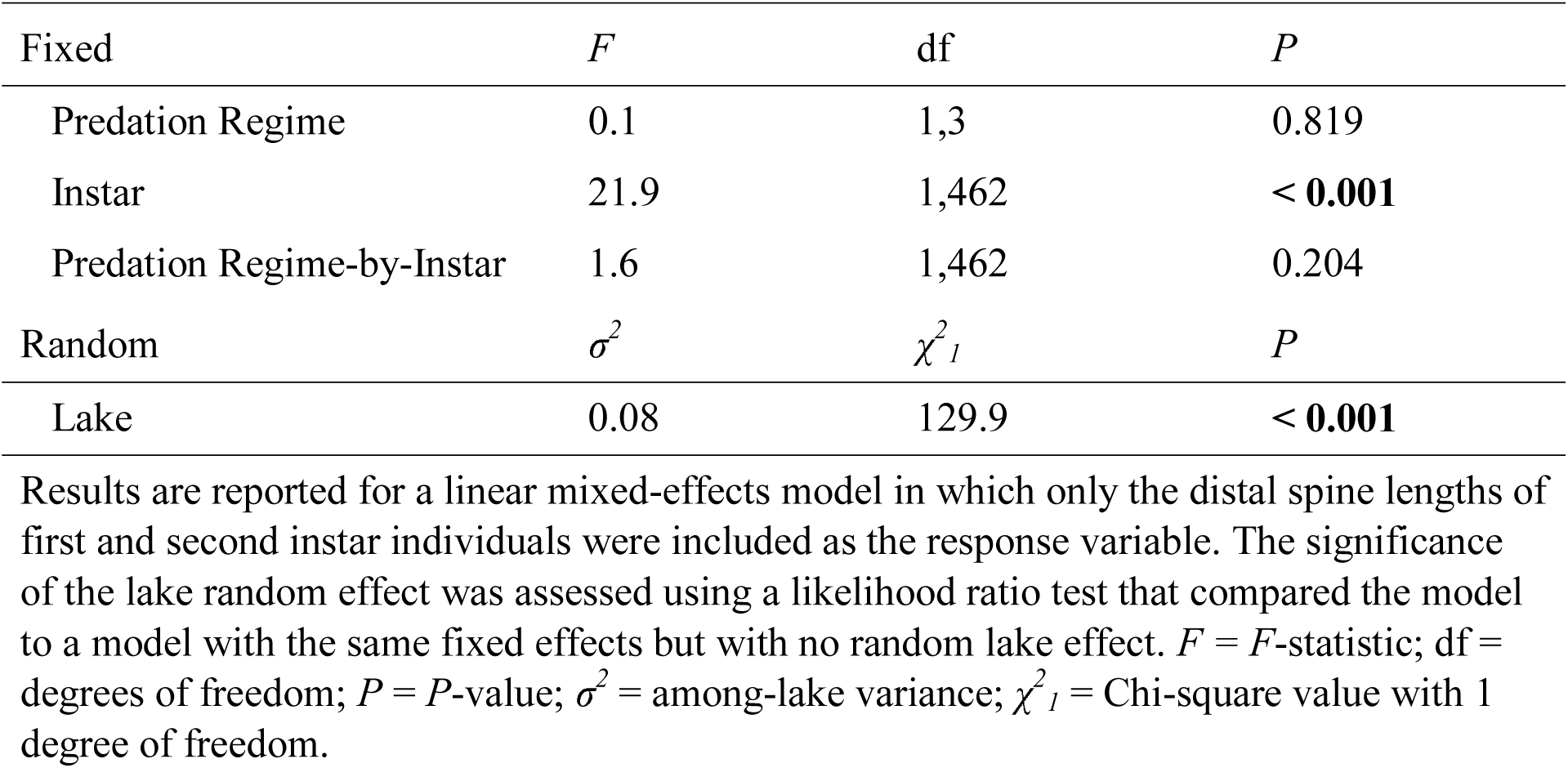
Natural selection on *Bythotrephes longimanus* distal spine length did not differ between predation regimes (i.e. GLP vs. NGLP).

### Common Garden Experiment

The mean distal spine length of second-generation individuals was 5.06 ± 0.04 mm (mean ± SE, SD = 0.39 mm), approximately 87% of the mean length observed in wild-caught individuals. Mean distal spine length of second-generation lab-born individuals did not differ among lakes (Table 5; Fig. 4). There was, however, significant genetic variation in *Bythotrephes* distal spine length, corresponding to a *H^2^* estimate of 0.24. Likewise, there was significant maternal variation in distal spine length, corresponding to a *m^2^* estimate of 0.61 (Table 6).

**Table 5.** The mean distal spine length of second-generation lab-born *Bythotrephes longimanus* did not differ among lakes.

| Fixed | $F$ | df | $P$ |
| --- | --- | --- | --- |
| Lake | 0.6 | 5,17 | 0.724 |
| Random | $\sigma^2$ | $\chi^2_1$ | $P$ |
| Subline nested within line | 0.10 | 62.2 | < **0.001** |
| Line | 0.05 | 22.2 | < **0.001** |

Results are reported for a linear mixed-effects model in which only the distal spine lengths of second-generation lab-born individuals were included as the response variable. The significance of the random effects was assessed using likelihood ratio tests that compared models with successively fewer random effects. $F$ = $F$ -statistic; df = degrees of freedom; $P$ = $P$ -value; $\sigma^2$ = variance; $\chi^2_1$ = Chi-square value with 1 degree of freedom.

**Fig. 4.**
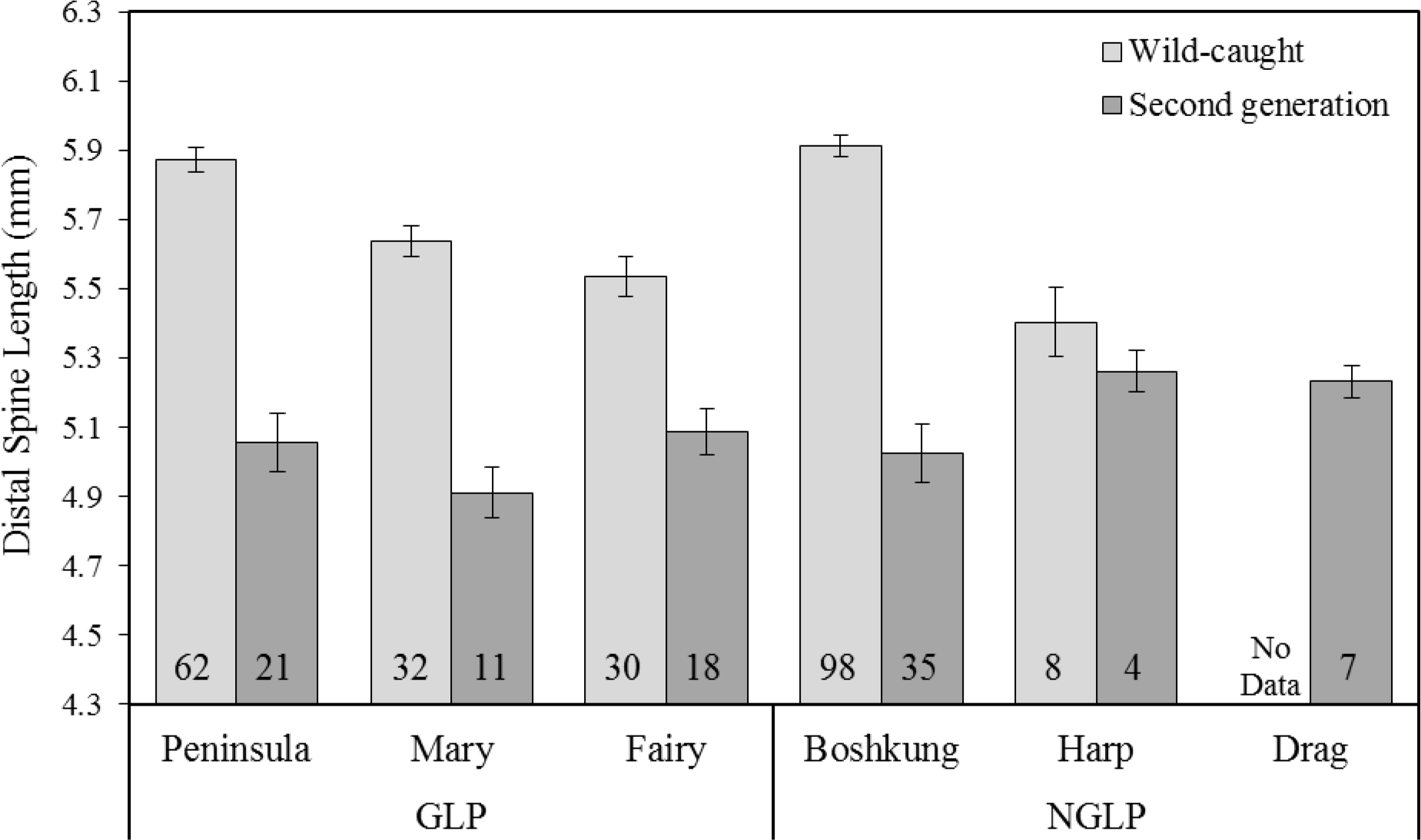
Mean distal spine length of wild-caught *Bythotrephes longimanus* and second-generation lab-born *Bythotrephes* for all study lakes. The mean distal spine length of second-generation lab-born individuals did not significantly differ among lakes (Table 5). Wild-caught animals were all first instar individuals so differences among lakes were not confounded by selection. The number in each bar represents the sample size for that lake. Error bars represent ± 1 standard error.

(Graphic created using Microsoft Excel 2013)

**Table 6.**
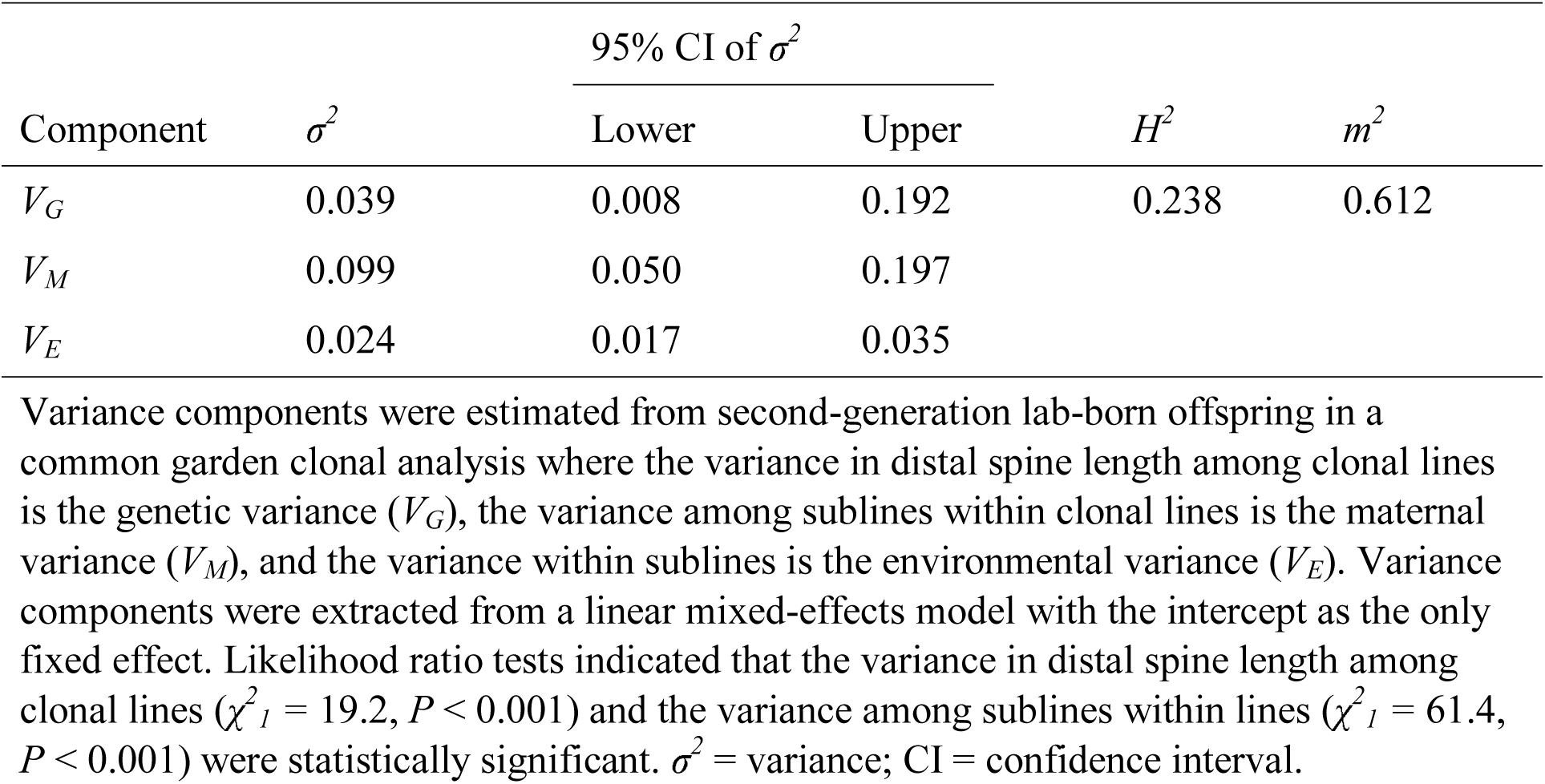
Genetic (*V_G_*), maternal (*V_M_*), and environmental (*V_E_*) variance components, broad-sense heritability (*H^2^*), and maternal effects (*m^2^*) for *Bythotrephes longimanus* distal spine length in Canadian Shield lakes from which *Bythotrephes* were sampled.

## Discussion

The goal of this study was to determine whether phenotypic differences in distal spine length among populations of *Bythotrephes* in Canadian Shield lakes could be explained by local adaptation or phenotypic plasticity. We found that *Bythotrephes* from two study lakes (Peninsula and Boshkung) had long mean distal spine lengths compared to those from three other study lakes (Mary, Fairy, and Harp); however, the mean distal spine lengths of second-generation lab-born individuals reared in a common environment did not differ among populations (Fig. 4).

Differences in distal spine length among populations were, therefore, a result of phenotypic plasticity in response to the maternal environment, and not local adaptation, despite spatially variable selection on this heritable trait. In view of *Bythotrephes* high dispersal capacity, the ability to plastically adjust distal spine length to local biotic conditions is consistent with the invasion success of this species over a relatively short time period.

The absence of genetically based differences in distal spine length among *Bythotrephes* populations was surprising because natural selection varied among populations. Specifically, we identified significant selection for longer distal spines in two of three GLP lakes (Mary and Fairy, but not Peninsula) and one of two NGLP lakes (Boshkung, but not Harp). A recent review of selection in wild populations identified that the median magnitude of directional selection (measured as the absolute value of standardized linear selection gradients) for survival was 0.08 (Kingsolver and Diamond 2011). Selection on *Bythotrephes* distal spine length that we measured in the three lakes with significant selection (Mary: *i* = 0.71; Fairy: *i* = 0.74; Boshkung: *i* = 0.82; Table 3) was thus very strong, falling within the top 10% of previously reported estimates (Kingsolver et al. 2001). Although Kingsolver and Diamond’s (2011) review reported selection using standardized selection gradients and we reported standardized selection differentials, these selection metrics have been found to often be similar in magnitude (Kingsolver and Diamond 2011). Miehls et al. (2014) also found strong selection for increased distal spine length in *Bythotrephes* in Mary Lake (*i* = 0.79) and Fairy Lake (*i* = 0.53) in the summer of 2008, suggesting that selection on *Bythotrephes* distal spine length in these lakes has been consistently strong.

Directional selection for increased distal spine length (which was observed in three of five lakes) should cause an evolutionary response if the trait is heritable (Falconer and Mackay 1996). We found significant genetic and maternal variation for *Bythotrephes* distal spine length, corresponding to a moderate broad-sense heritability and large maternal effect (Table 6; Mousseau and Roff 1987), which are the first such estimates for *Bythotrephes* in Canadian Shield lakes. Our estimates of genetic variation and heritability for *Bythotrephes* distal spine length were very similar to previous estimates in Lake Michigan in July (*V_G_* = 0.06, *H*^2^ = 0.27; Miehls et al. 2012), indicating that most of the genetic variation in distal spine length has been maintained since *Bythotrephes* invasion from the Laurentian Great Lakes, and that the spread of *Bythotrephes* has not limited their potential for adaptive evolution. Despite this adaptive potential, and significant differences in selection among lakes, we found no evidence of genetic differentiation for distal spine length among populations of *Bythotrephes*. Previous work on *Bythotrephes* used historic and contemporary wild-caught animals and remnant distal spines retrieved from sediment cores to test for a response to selection on distal spine length since *Bythotrephes* invasion of Lake Michigan, and found little evidence of phenotypic change through time (Miehls et al. 2015). Together, our results and those of Miehls et al. (2015) provide clear examples of selection on a heritable trait not leading to an evolutionary response temporally (in Lake Michigan) or spatially (in these Canadian Shield lakes).

There are several reasons why selection on a heritable trait may not cause evolutionary change (i.e. evolutionary stasis; Merilä et al. 2001). For example, temporal fluctuations in selection can influence the direction and strength of selection overall (Siepielski et al. 2009; Bell 2010; Kingsolver and Diamond 2011). In GLP lakes, predation risk for *Bythotrephes* increases through the growing season because juvenile gape-limited fish grow from sizes too small to consume any *Bythotrephes* to sizes that can consume some *Bythotrephes* depending on gape-size (Straile and Halbich 2000; Branstrator 2005; Pothoven et al 2012; Miehls et al. 2015). Our study only looked at a single snapshot of selection for each study lake, but previous work on *Bythotrephes* in Lake Michigan found strong temporal variation in selection within a growing season, which reduced net selection (Miehls et al. 2015). Selection might also fluctuate across years. Miehls et al. (2014) found significant selection for increased distal spine length in Peninsula, Mary, and Fairy (GLP lakes; *i* = 0.20-0.79) in 2008, but no selection in Boshkung and Harp (NGLP lakes) in 2008, whereas we found significant selection in Mary, Fairy, and Boshkung (*i* = 0.71-0.82; Table 3), but no selection in Peninsula and Harp in 2014.

It is also possible that *Bythotrephes* experience a tradeoff between components of fitness (i.e. survival vs. fecundity; Roff 2002) or that selection varies among life stages (Schluter et al. 1991) such that our selection estimates based on a fitness component (i.e. survival between first and second instar) does not represent overall selection. Previous work suggested that *Bythotrephes* exhibit a tradeoff between clutch size and offspring distal spine length (i.e. females that produced offspring with longer distal spines had smaller clutches; Straile and Halbich 2000; Pothoven et al. 2003; Miehls et al. 2013), which means that viability selection favouring longer distal spines could be opposed by fecundity selection favouring shorter distal spines. Lastly, we measured selection between first and second instar individuals, but selection on later instar stages is likely to occur on the length of the total spine, rather than just the distal spine (Fig. 1), which might reduce the strength of selection on distal spine length if selection on the total spine is weaker or in the opposite direction (see Miehls et al. 2015 for further discussion of potential causes of stasis in *Bythotrephes*).

In contrast with previous findings (i.e. Miehls et al. 2014), the differences among lakes in mean distal spine length and selection that we observed were inconsistent with GLP as an agent of selection. In particular, our results from two lakes did not match our expectations. First, the population in Peninsula Lake (GLP) had a long mean distal spine length, but weak selection (Table 3; Fig. 3). It is possible that individuals from Peninsula did not experience selection because distal spine lengths were already long enough to provide defense against GLP. Comparing GLP lakes, individuals from Fairy had the shortest mean distal spines (5.54 mm) but the strongest selection (*i* = 0.74), whereas individuals from Peninsula had the longest mean distal spines (5.87 mm) but the weakest selection (*i* = 0.03; Fig. 3; see also Miehls et al. 2014). These results suggest the possibility of a threshold distal spine length that provides refuge from GLP, which is consistent with “hard” natural selection (Wallace 1975).

The second unexpected finding was that the *Bythotrephes* in Boshkung Lake had long distal spines and experienced strong selection, despite Boshkung being classified as a NGLP lake (Table 3; Fig. 3). The most obvious explanation for this finding is that Boshkung may no longer be dominated by NGLP. Rainbow smelt (the dominant gape-limited fish predator) were previously found to be present in Boshkung (Young and Yan 2008), but cisco (the dominant non-gape-limited fish predator) were thought to be the dominant predator of *Bythotrephes* (Table 1; Strecker et al. 2006; Miehls et al. 2014). Unfortunately, there has not been a recent fish survey of Boshkung to provide further insights into whether the strong selection measured in this study could be explained by an increase in the abundance of smelt relative to cisco since the last survey. Interestingly, Miehls et al. (2014) also found that the *Bythotrephes* population in Boshkung had a longer mean distal spine length compared to Harp (a NGLP lake with no smelt), but their primary analysis yielded no evidence of selection. However, an alternative analysis found the occurrence of reasonably strong (though statistically insignificant) selection (*i* = 0.46) that was stronger than selection in Peninsula (GLP) (*i* = 0.32), which is somewhat consistent with our findings. More recent fish surveys in Boshkung Lake will be needed to determine whether Boshkung is in fact a GLP lake, and more replication of NGLP and GLP lakes will be needed to assess the importance of gape-limited predation as an agent of selection on *Bythotrephes* distal tail spine length.

It is also possible that natural selection is affected not only by the presence of gape-limited predators, but also by the behavioural exposure of *Bythotrephes* to gape-limited predators. While adult rainbow smelt are typically in the hypolimnion (Evans and Loftus 1987), gape-limited predation of *Bythotrephes* by smelt is due to smaller size classes of smelt (Barnhisel and Harvey 1995) that are often in shallower water (Evans and Loftus 1987). Diel vertical migration provides a possible refuge from visual predators in the epilimnion during the daytime, which might also affect exposure to gape-limited smelt and hence natural selection on distal spine length. Young and Yan (2008) found that lakes containing cisco reduced diel vertical migration of *Bythotrephes* compared to lakes without cisco. Increased diel vertical migration in lakes without cisco (i.e. GLP lakes) might, therefore, have weakened natural selection on distal spine length from gape-limited predation below what it might otherwise be in the absence of this behavioural refuge. In contrast, reduced diel vertical migration in response to cisco might have enhanced the exposure of *Bythotrephes* to the small number of gape-limited smelt reported to be in Boshkung Lake, which could provide a possible explanation for the stronger than expected natural selection in that lake. The degree to which our estimates of natural selection were affected by diel vertical migration cannot be known, but it is clear that natural selection on *Bythotrephes* distal spine length will depend on both the presence and behavioural exposure to gape-limited predators.

Overall, our results strongly support the hypothesis that phenotypic differences among *Bythotrephes* populations in these study lakes are a result of plasticity, but the way in which plasticity causes these differences remains unclear. In general, there are two ways in which plasticity can result in the phenotypic differences among populations that we observed (Fig. 3). First, it is possible that the reaction norm for distal spine length is the same in all populations, and that phenotypic differences among populations are a result of differences in the level of an environmental cue among lakes (Fig. 5a). Alternatively, phenotypic differences among populations of *Bythotrephes* could have resulted from the evolution of different reaction norm slopes among lakes in response to spatial variation in selection (Fig. 5b).

**Fig. 5.**
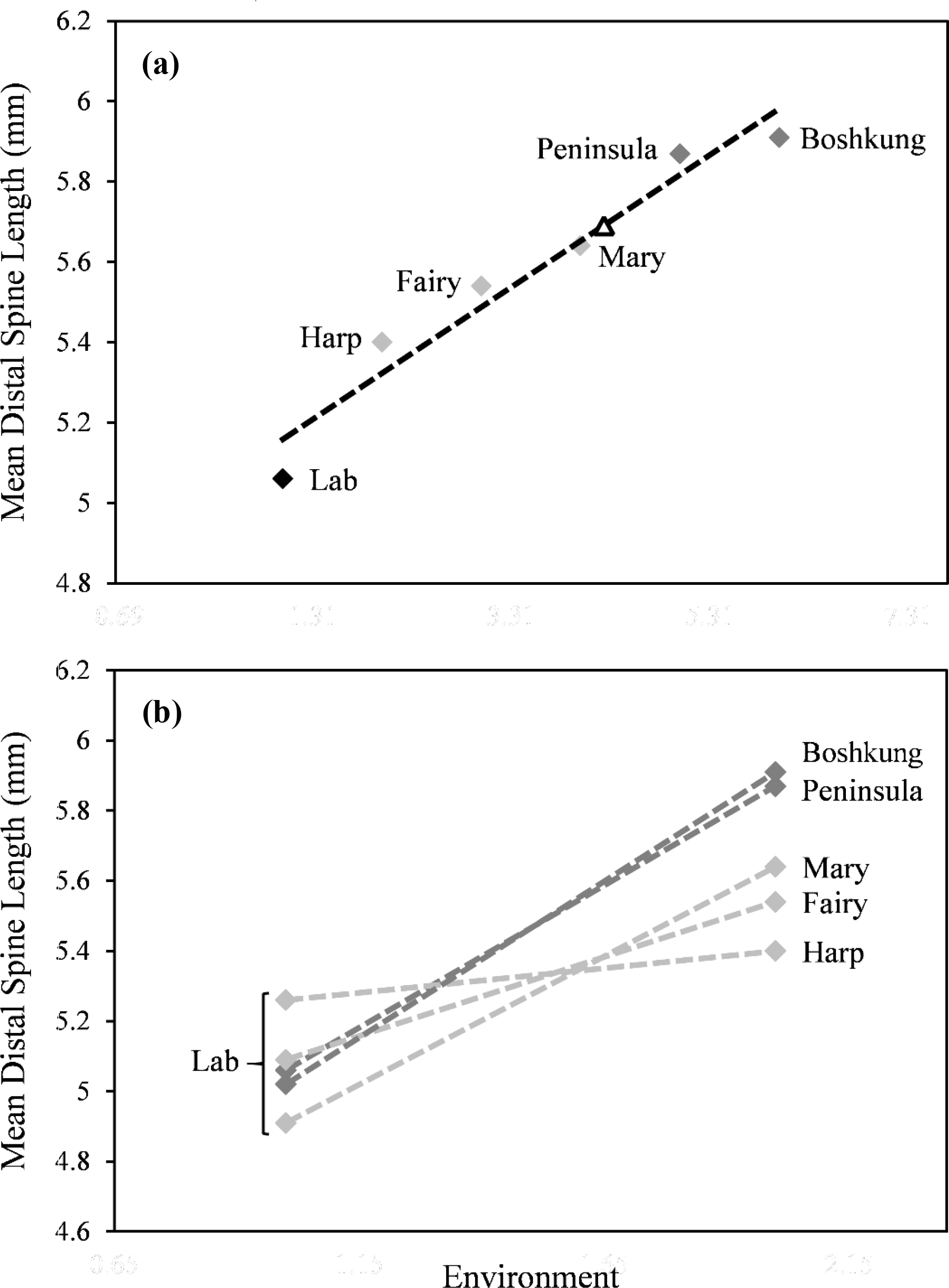
(a) Hypothetical reaction norm for *Bythotrephes longimanus* in Canadian Shield lakes if the reaction norm for distal spine length is the same in all study populations. The “lab” value is the mean distal spine length for second-generation lab-born individuals pooled across lakes. The white triangle represents the mean phenotype that would be expected for lab-reared *Bythotrephes* if the environmental cue for plasticity was present in the common garden medium. (b) Hypothetical reaction norms for *Bythotrephes longimanus* populations from each study lake if populations evolved different reaction norm slopes in response to spatial variation in selection.

(Graphic created using Microsoft Excel 2013)

The environmental cue that caused the plastic response that we observed is also unclear. Our finding that the mean distal spine length of common garden *Bythotrephes* was smaller than that of wild caught *Bythotrephes* for all lakes (Fig. 4) suggests that the cue was not present in the water that we collected to create the common garden medium. Had the cue been present in the common medium, we would have expected the phenotypes from lab-reared *Bythotrephes* to converge on an intermediate phenotype (Fig. 5a), or to maintain phenotypic differences among lake populations as observed in wild-caught animals (Fig. 5b). Miehls et al. (2013) found that *Bythotrephes* from Lake Michigan induce longer tail spines in their offspring in response to warmer water temperature, and the water temperature in our common environment (20°C) was slightly lower than lake temperatures at the time of collection (range = 20.5°C to 21.7°C; Table 1). However, the magnitude of the differences in water temperature among lakes were quite small (Table 1), and these temperature differences were inconsistent with phenotypic differences in distal spine length among lakes. For example, among GLP lakes, Fairy and Mary had shorter mean distal spine lengths than Peninsula, but water temperature was cooler in Mary (21.2°C) and warmer in Fairy (21.7°C) compared to Peninsula (21.5°C). Miehls et al. (2013) also found that *Bythotrephes* distal spine length did not change in response to kairomones from yellow perch (*Perca flavescens*), so it is unlikely that differences in the presence or concentration of fish kairomones among Canadian Shield lakes caused the observed phenotypic differences. It is possible that *Bythotrephes* respond to kairomones of specific fish species (as opposed to fish kairomones in general) and that this cue somehow degraded in the lab, but this cue would need to be specific to gape-limited predators for which the induction of longer distal spines would be beneficial, and not a generic cue of predation risk (Miehls et al. 2013). Clearly, further work is needed to identify the causes of phenotypic differences in distal spine lengths among lakes. This would involve identifying the environmental variable causing plastic responses in distal spine lengths as well as the degree to which this plasticity varies among lakes.

In conclusion, we have demonstrated that phenotypic differences in a key trait involved in interspecific interactions are a result of phenotypic plasticity and not local adaptation, despite spatially variable selection on a heritable trait. This evolutionary stasis (Merilä et al. 2001) serves as a reminder that adaptation cannot be inferred from phenotypic divergence even when this divergence is consistent with natural selection. Furthermore, this finding highlights the potential importance of phenotypic plasticity as a mechanism by which invasive species might respond to environmental heterogeneity (e.g. Dybdahl and Kane 2005). General lessons about the importance of local adaptation and phenotypic plasticity to the spread of exotic species, however, require further explicit tests of their relative importance across a wider range of taxa.

## Acknowledgements

We thank Teresa Crease and Beren Robinson for insightful comments, and for providing field and lab equipment. The Dorset Environmental Science Centre (DESC) provided a climate controlled facility. James Rusak (DESC) and Matt Cornish (Hagen Aqualab) assisted with lab set-up. Andrea Miehls helped with *Bythotrephes* collection and culturing protocols. Guang Zhang assisted with field work. Emily De Freitas, Evan McKenzie, Kaileigh Watson, Katelyn Cross, Kirsten Bradford, Marissa Skinner, Mary Paquet, Meera Navaratnam, Ronena Wolach, and Yu Jin Song assisted with data collection. This work was funded by an Ontario Ministry of Research and Innovation Early Researcher Award to Andrew McAdam, a Natural Sciences and Engineering Research Council Discovery Grant to Andrew McAdam, and an Ontario Graduate Scholarship to Giuseppe Fiorino.

## Literature Cited

Abdi H, Williams LJ (2010) Tukey’s honestly significant difference (HSD) test. In: Salkind NJ (ed) Encyclopedia of Research Methods. Sage, Thousand Oaks, CA.

Abramoff MD, Magelhaes PJ, Ram SJ (2004) Image processing with ImageJ. Biophotonics International 11:36–42.

Allendorf FW, Lundquist LL (2003) Introduction: population biology, evolution, and control of invasive species. Conserv Biol 17:24–30.

Barnhisel, DR, Harvey HA (1995) Size-specific fish avoidance of the spined crustacean *Bythotrephes*: field support for laboratory predictions. Can J Fish Aquat Sci 52:768–775.

Barnhisel DR (1991a) The caudal appendage of the cladoceran *Bythotrephes cederstroemi* as defense against young fish. J Plankton Res 13:529–537.

Barnhisel DR (1991b) Zooplankton spine induces aversion in small fish predators. Oecologia 88:444–450.

Bell G (2010) Fluctuating selection: the perpetual renewal of adaptation in variable environments. Philos Trans R Soc Lond B Biol Sci 365:87–97.

Branstrator DK (2005) Contrasting life histories of the predatory cladocerans *Leptodora kindtii* and *Bythotrephes longimanus*. J Plankton Res 27:569–585.

Bunnell DB, Davis BM, Warner DM, Chriscinske MA, Roseman EF (2011) Planktivory in the changing Lake Huron zooplankton community: *Bythotrephes* consumption exceeds that of *Mysis* and fish. Freshwater Biol 56:1281–1296.

Burkhardt S (1994) Seasonal size variation in the predatory cladoceran *Bythotrephes cederstroemii* in Lake Michigan. Freshwater Biol 31:97–108.

Colautti RI, Manca M, Viljanen M, Ketelaars HAM, Bürgi H, Macisaac HJ, Heath DD (2005) Invasion genetics of the Eurasian spiny waterflea: evidence for bottlenecks and gene flow using microsatellites. Mol Ecol 14:1869–1879.

Day T, Abrams PA, Chase JM (2002) The role of size-specific predation in the evolution and diversification of prey life histories. Evolution 56:877–887.

Drake JA, Mooney HA, di Castri F, Groves RH, Kruger FJ, Rejmanek M, Williamson M (1989) Biological Invasions: A Global Perspective. John Wiley and Sons, New York, NY.

Dybdahl MF, Kane SL (2005) Adaptation vs. phenotypic plasticity in the success of a clonal invader. Ecology 86:1592–1601.

Dzialowski AR, Lennon JT, O’Brien WJ, Smith VH (2003) Predator-induced phenotypic plasticity in the exotic cladoceran *Daphnia lumholtzi*. Freshwater Biol 48:1593–1602.

Evans DO, Loftus DH (1987) Colonization of inland lakes in the Great Lakes region by rainbow smelt, *Osmerus mordax*: their freshwater niche and effects on indigenous fishes. Can J Fish Aquat Sci 44:249–266.

Facon B, Genton BJ, Shykoff J, Jarne P, Estoup A, David P (2006) A general eco-evolutionary framework for understanding bioinvasions. Trends Ecol Evol 21:130–135.

Falconer DS, Mackay TFC (1996) Introduction to Quantitative Genetics (4th ed). Longman, Harlow, UK.

Godoy O, Saldana A, Fuentes N, Valladares F, Gianoli E (2011) Forests are not immune to plant invasions: Phenotypic plasticity and local adaptation allow *Prunella vulgaris* to colonize a temperate evergreen rainforest. Biol Invasions 13:1615–1625.

Grigorovich IA., Pashkova OV, Gromova YF, van Overdijk CDA (1998) *Bythotrephes longimanus* in the Commonwealth of Independent States: variability, distribution and ecology. Hydrobiologia 379:183–198.

Hovius JT, Beisner BE, McCann KS (2006) Epilimnetic rotifer community responses to *Bythotrephes longimanus* invasion in Canadian Shield lakes. Limnol Oceanogr 51:1004–1012.

Johannsson OE, Mills EL, O’Gorman R (1991) Changes in the nearshore and offshore zooplankton communities in Lake Ontario: 1981-88. Can J Fish Aquat Sci 48:1546–1557.

Kawecki TJ, Ebert D (2004) Conceptual issues in local adaptation. Ecol Lett 7:1225–1241.

Kelly NE, Yan ND, Walseng B, Hessen DO (2013) Differential short- and long-term effects of an invertebrate predator on zooplankton communities in invaded and native lakes. Divers Distrib 19:396–410.

Kim N, Yan ND (2013) Food limitation impacts life history of the predatory cladoceran *Bythotrephes longimanus*, an invader to North America. Hydrobiologia 715:213–224.

Kim N, Yan ND (2010) Methods for rearing the invasive zooplankter *Bythotrephes* in the laboratory. Limnol Oceanogr: Meth 8:552–561.

Kingsolver JG, Diamond SE (2011) Phenotypic selection in natural populations: what limits directional selection? Am Nat 177:346–357.

Kingsolver JG, Hoekstra HE, Hoekstra JM, Berrigan D, Vignieri SN, Hill CE, Hoang A, Gilbert P, Beerli P (2001) The strength of phenotypic selection in natural populations. Am Nat 157:245–261.

Lambrinos JG (2004) How interactions between ecology and evolution influence contemporary invasion dynamics. Ecology 85:2061–2070.

Lee CE (2002) Evolutionary genetics of invasive species. Trends Ecol Evol 17:386–391.

Lockwood JL, Cassey P, Blackburn T (2005) The role of propagule pressure in explaining species invasions. Trends Ecol Evol 20:223–228.

Lüning J (1992) Phenotypic plasticity of *Daphnia pulex* in the presence of invertebrate predators: morphological and life history responses. Oecologia 92:383–390.

Lynch M, Walsh B (1998) Genetics and Analysis of Quantitative Traits. Sinauer, Sunderland, MA.

Mack RN, Simberloff D, Lonsdale WM, Evans H, Clout M, Bazzaz FA (2000) Biotic invasions: causes, epidemiology, global consequences, and control. Ecol Appl 10:689–710.

Merilä J, Sheldon BC, Kruuk LEB (2001) Explaining stasis: Microevolutionary studies in natural populations. Genetica 112-113:199–222.

Miehls ALJ, McAdam AG, Bourdeau PE, Peacor SD (2013) Plastic response to a proxy cue of predation risk when direct cues are unreliable. Ecology 94:2237–2248.

Miehls ALJ, Peacor SD, McAdam AG (2014) Gape-Limited Predators As Agents of Selection on the Defensive Morphology of an Invasive Invertebrate. Evolution 68:2633–2643.

Miehls ALJ, Peacor SD, McAdam AG (2012) Genetic and maternal effects on tail spine and body length in the invasive spiny water flea *(Bythotrephes longimanus)*. Evol Appl 5:306–316.

Miehls ALJ, Peacor SD, Valliant L, McAdam AG (2015) Evolutionary stasis despite selection on a heritable trait in an invasive zooplankton. J Evol Biol 28:1091–1102.

Mooney HA, Cleland EE (2001) The evolutionary impact of invasive species. Proc Natl Acad Sci USA 98:5446–5451.

Mousseau TA, Fox CW (1998) The adaptive significance of maternal effects. Trends Ecol Evol 13:403–407.

Mousseau TA, Roff DA (1987) Natural selection and the heritability of fitness components. Heredity 59:181–197.

Novak SJ (2007) The role of evolution in the invasion process. Proc Natl Acad Sci USA 104:3671–3672.

Parker IM, Rodriguez J, Loik ME (2003) An evolutionary approach to understanding the biology of invasions: local adaptation and general-purpose genotypes in the weed *Verbascum thapsus*. Conserv Biol 17:59–72.

Pfennig DW, Wund MA, Snell-Rood EC, Cruickshank T, Schlichting CD, Moczek AP (2010) Phenotypic plasticity’s impacts on diversification and speciation. Trends Ecol Evol 25:459–467.

Pigliucci M (2005) Evolution of phenotypic plasticity: where are we going now? Trends Ecol Evol 20:481–486.

Pinheiro JC, Bates DM (2000) Mixed-Effects Models in S and S-PLUS. Springer-Verlag, New York, NY.

Pinheiro JC, Bates DM, DebRoy S, Sarkar D, R Core Team (2015) nlme: Linear and Nonlinear Mixed Effects Models. R package version 3.1-121.

Pothoven SA, Fahnenstiel GL, Vanderploeg HA (2003) Population characteristics of *Bythotrephes* in Lake Michigan. J Great Lakes Res 29:145–156.

Pothoven SA, Vanderploeg HA, Warner DM, Schaeffer JS, Ludsin SA, Claramunt RM, Nalepa TF (2012) Influences on *Bythotrephes longimanus* life-history characteristics in the Great Lakes. J Great Lakes Res 38:134–141.

Pothoven SA, Vanderploeg HA, Cavaletto JF, Krueger DM, Mason DM, Brandt SB (2007) Alewife planktivory controls the abundance of two invasive predatory cladocerans in Lake Michigan. Freshwater Biol 52:561–573.

R Core Team (2015) R: A language and environment for statistical computing. R Foundation for Statistical Computing, Vienna, Austria.

Roff D (2002) Life History Evolution. Sinauer Associates, Sunderland, MA.

Shea K, Chesson P (2002) Community ecology theory as a framework for biological invasions. Trends Ecol Evol 17:170–176.

Schluter D, Price TD, Rowe L (1991) Conflicting selection pressures and life history trade-offs. Proc R Soc London Ser B Bio 246:11–17.

Si C-C, Dai Z-C, Lin Y, Qi S-S, Huang P, Miao S-L, Du D-L (2014) Local adaptation and phenotypic plasticity both occurred in *Wedelia trilobata* invasion across a tropical island. Biol Invasions 16:2323–2337.

Siepielski AM, DiBattista JD, Carlson SM (2009) It’s about time: the temporal dynamics of phenotypic selection in the wild. Ecol Lett 12:1261–1276.

Straile D, Halbich A (2000) Life History and Multiple Antipredator Defenses of an Invertebrate Pelagic Predator, *Bythotrephes longimanus*. Ecology 81:150–163.

Strecker AL, Arnott SE, Yan ND, Girard R (2006) Variation in the response of crustacean zooplankton species richness and composition to the invasive predator *Bythotrephes longimanus*. Can J Fish Aquat Sci 63:2126–2136.

Therriault TW, Grigorovich IA, Cristescu ME, Ketelaars HAM, Viljanen M, Heath DD, Macisaac HJ (2002) Taxonomic resolution of the genus *Bythotrephes* Leydig using molecular markers and re-evaluation of its global distribution. Divers Distrib 8:67–84.

Urban MC (2007) Predator size and phenology shape prey survival in temporary ponds. Oecologia 154:571–580.

Urban MC (2008) Salamander evolution across a latitudinal cline in gape limited predation risk. Oikos 117:1037–1049.

Vitousek PM, D’Antonio CM, Loope LL, Westbrooks R (1996) Biological Invasions as Global Environmental Change. Am Sci 84:468–478.

Wallace B (1975) Hard and Soft Selection Revisited. Evolution 29:465–473.

West-Eberhard MJ (2003) Developmental Plasticity and Evolution. Oxford University Press, Oxford, UK.

Young JD, Yan ND (2008) Modification of the diel vertical migration of *Bythotrephes longimanus* by the cold-water planktivore, *Coregonus artedi*. Freshwater Biol 53:981–995.

Yurista PM (1992) Embryonic and postembryonic development in *Bythotrephes cederstoemii*. Can J Fish Aquat Sci 49:1118–1125.

